# Premarin has opposing effects on spatial learning, neural activiation and serum cytokine levels in middle age dependent on reproductive history

**DOI:** 10.1101/234229

**Authors:** Liisa AM Galea, Meighen M. Roes, Christina van den Brink, Carmen Chow, Rand Mahmoud, Stephanie E. Lieblich, Paula Duarte-Guterman

## Abstract

Menopause is associated with cognitive decline, and hormone therapies (HT) can improve cognition dependent on time since menopause. Previous parity also influences cognition in later life. The present study investigated how primiparity and long-term ovariectomy influence cognition, hippocampal neurogenesis, and neuronal activation in middle-aged rats in response to the HT, Premarin. Nulliparous and primiparous rats were sham-ovariectomized or ovariectomized, administered vehicle or Premarin six months later, and trained in the Morris water maze. Premarin improved spatial learning and memory in nulliparous rats, but impaired spatial and reversal learning in primiparous rats. Primiparity increased hippocampal neurogenesis, whereas Premarin treatment decreased immature neurons in both primiparous and nulliparous middle-aged rats in a region-specific manner. Moreover, Premarin increased serum TNFα and KC/GRO in nulliparous, but not primiparous, rats, whereas Premarin increased zif268 expression in the CA3 region of the hippocampus in primiparous rats. Thus, primiparity alters how Premarin affects spatial learning, neuronal activation, serum cytokines, and adrenal mass. These findings have implications for the tailored treatment of age-associated cognitive decline in women.

---

Maternal adaptation describes how the women’s body has to adapt to allow for fetal growth as a host of physiological changes must occur to allow for appropriate survival of the fetus. For example, cardiac output, and pulmonary function are increased or decreased by as much as 50% in gestating women (Grindheim et al., 2012; Savu et al., 2012). Furthermore, the endocrine system is modified, as the placenta releases a variety of hormones in high concentrations such as estradiol, progesterone, and CRH (Brett and Baxendale, 2001; Holl et al., 2009). The mother’s immune system also undergoes modifications during pregnancy in part to foster tolerance to the forming fetus (Ghaebi et al., 2017). Although it has been widely assumed that many aspects of maternal physiology normalize after the expulsion of the placenta, certain physiological changes outlast the pregnancy and early postpartum or even emerge later in life. For example, as parous women age, mid-luteal phase estradiol levels are decreased compared to nulliparous women (Dorgan et al., 1995). In addition, parity alters the immune profile of aged animals (Barrat et al., 1997; Mahmoud et al., under revision). Parity also has lasting consequences on the brain, as long-lasting changes in hippocampal volume and cognition are seen in women and rodents (for review see Roes & Galea, 2015; Hoekzmsa et al., 2017; Galea et al., 2000; Galea et al., 2014; Barha and Galea, 2015). Primiparas women exhibit reductions of hippocampal volume, and in other brain regions, two months after parturition that still are evident 2 years post parturition (Hoekzmsa et al., 2017). In addition, primiparous rats show reduced hippocampal volume and neurogenesis during lactation and throughout the postpartum period (Galea et al., 2000; Pawluski and Galea, 2007, Leuner and Gould, 2007; Workman et al., 2015). Thus, studies suggest that reproductive experience has lasting effects on the maternal physiology and hippocampal structure and plasticity.

Neurogenesis in the hippocampus is reduced in the early and late postpartum in primiparous female rats (Galea et al., 2000; Pawluski and Galea, 2007, Leuner and Gould, 2007; Workman et al., 2015) but curiously in middle age, increased hippocampal neurogenesis is seen in multiparous rats relative to nulliparous rats (Barha et al., 2015). Multiparous rodents have more immature neurons, increased levels of brain derived neurotrophic factor (BDNF), and synaptophysin and spinophilin than nulliparous rats in middle age (Barha et al., 2015; Cui et al. 2014; Macbeth et al., 2008; Rosetti et al, 2016). Furthermore, in response to estrogens, multiparous middle-aged rats show an enhancement in cell proliferation whereas nulliparous middle-aged rats do not, a pattern seen in younger adult female rats. In the hippocampus and amygdala, neuropathology (neuritic plaques and neurofibrillary tangles) was positively correlated with parity in older women (Beeri et al., 2009), suggesting some neurological consequences of increased parity in women. These findings suggest that increased parity is associated with both increased hippocampal plasticity and pathology well into middle age, long after offspring have been reared.

Previous parity also influences cognitive ability in middle- and older-aged rodents and perhaps women. Primiparous and multiparous middle-aged rats display better performance compared to nulliparous rats in the dry land maze (Gatewood et al., 2005; Love et al., 2005), water maze (Lemaire et al., 2006; Barha et al., 2015) and in reversal learning tasks (Gatewood et al., 2005). Importantly, previous reproductive experience may affect various aspects of hippocampal cognition differently. In middle age, multiparous rats have enhanced early acquisition in the spatial working memory version of the Morris water maze, but impaired reference memory acquisition, compared to nulliparous rats (Barha et al, 2015). Studies in women are more equivocal. However, parity is associated with better memory in older women with factors such as age of first pregnancy, genotype, and amount of parity playing a moderating role (Fox et al., 2013; Karim et al., 2016; Roes and Galea, 2015). Taken together these studies suggest that previous parity may be associated with improvements in learning and memory in middle-age.

Menopause is associated with a decline in circulating estrogens as well as declines in certain cognitive domains (Weber, Rubin & Maki, 2013). Conversely, hormone therapy (HT) in postmenopausal women can improve cognition depending on age at administration and HT composition (Hogeverst et al., 2000; Ryan et al., 2008). Premarin is a common HT of conjugated equine estrogens (CEE), composed 50% of estrone sulphate and 0.1% estradiol sulphate. Metaanalyses indicate that fewer studies report positive effects on cognition with Premarin compared to estradiol-based therapies (Hogevorst et al., 2000; Ryan et al., 2008). Furthermore, the effects of HT on cognition in women were dependent on the timing of when HTs were prescribed (early or late in menopause), with a greater proportion of studies finding detrimental outcomes, or fewer positive outcomes, when HT is initiated 10 or more years after menopause. In animal studies, a ‘critical window’ exists during which estradiol administration early after ovariectomy (OVX) improves hippocampal cognition in middle age, but estradiol has no effects on hippocampal cognition after long-term OVX (Walf, Paris & Frye, 2009; Gibbs 2010). However, duration of ovariectomy itself can influence cognition, with long-term (6 months) OVX improving spatial memory in nulliparous rats (Bimonte-Nelson et al 2003). Intriguingly, the study showing beneficial effects of long-term ovariectomy have used nulliparous rats (Bimonte-Nelson et al., 2003), whereas one study showing no cognitive benefit of estradiol after long-term ovariectomy used multiparous rats (Walf, Paris & Frye, 2009), suggesting that parity modulates the ovariectomy-induced effects on cognition in later life.

Reproductive experience alters the cognitive and neuroplastic effects of estrogens. Estrone and estradiol administration upregulate cell proliferation in the hippocampus of middle-aged multiparous but not nulliparous rats (Barha & Galea, 2011), suggesting that parity preserves the sensitivity of the hippocampus to estradiol and estrone later in life. Furthermore, ovariectomy in middle age has opposing effects on memory in nulliparous versus multiparous rats, improving spatial memory retrieval in nulliparous rats but impairing it in multiparous rats (Barha et al., 2015). Thus, reproductive experience alters the hippocampus-dependent memory and neuroplastic response to estrogens in middle age. The mechanisms by which parity can alter the aging trajectory are unclear, but may involve changes in inflammatory signatures, (Mahmoud et al., under review; Barrat et al., 1997; Cramer et al., 2017), or levels of estrogens (Dorgan et al., 1995; Bridges and Byrnes, 2006; Barrett et al., 2014), and both of these factors were examined in the present study.

The aim of the present study was to determine the effects of primiparity, Premarin, and long-term ovariectomy on spatial memory, neurogenesis and neuronal activation in the hippocampus of middle-aged rats. We also examined possible mediating factors such as adrenal mass, serum levels of cytokines, and estrogens. We hypothesized that Premarin treatment in middle-aged rats would affect spatial reference and reversal learning, neurogenesis, and neuron activation in the hippocampus in a manner that is dependent on reproductive experience and hormone status.

## Method

### Animals

Ninety-seven female and 16 male Sprague-Dawley rats (Charles River, Quebec) were 2 months old at arrival at the University of British Columbia. Rats were housed in opaque polyurethane bins (24 x 16 x 46 cm) with aspen chip bedding, and were given standard laboratory chow (Harlan, Canada) and tap water ***ad libitum.*** Rats were maintained under a 12h:12h light/dark cycle (lights on at 07:00h). Rats were randomly assigned to parity (primiparous (Prim) or nulliparous (Null)), surgery (ovariectomy (OVX) or sham surgery), and treatment (oil or Premarin) conditions (*n*’ s = 12-13 before attrition). Thus, this resulted in 8 groups: Null-Sham-Oil; Null-Sham-Prem; Null-OVX-Oil; Null-OVX-Prem; Prim-Sham-Oil; Prim-Sham-Prem; Prim-OVX-Oil; Prim-OVX-Prem.

Rats were double-housed except during pregnancy and gestation (nulliparous females were single-housed for an equivalent period of time) and recuperation from surgery. To prevent nulliparous rats from being exposed to odors and noises from males and/or pre-weaning pups, nulliparous rats were housed in a separate colony room until breeding and weaning were complete. All testing was conducted in accordance with ethical guidelines set by the Canada Council for Animal Care and all procedures were approved by the University of British Columbia Animal Care Committee. All efforts were made to reduce the number and suffering of animals.

### Apparatus

The Morris Water Maze was a black circular pool, 180cm in diameter, filled with room temperature water mixed with black tempura paint (non-toxic) to render it opaque. Large and distinct distal cues were placed on all four walls of the room surrounding the pool and remained constant throughout behavioral testing. In the water maze task, the animal must use the distal cues to navigate to a hidden platform, submerged roughly 2 cm beneath the pool surface. Cameras installed above the center of the pool were connected to ANY-maze 4.98 software (Stoelting, Wood Dale, IL, USA) in order to record measures.

### Breeding and procedures

See Figure 1 for experimental timeline. During breeding, two females and one male were paired overnight beginning at approximately 5pm. Females were vaginally lavaged each morning between 7:30 and 9:00 am and samples were assessed for the presence of sperm. Upon identification of sperm, females were considered pregnant, weighed, and single housed into clean cages. One day after birth, all litters were culled to 5 males and 5 females. If there were not enough males or females in one litter, pups were cross-fostered from a dam that gave birth the same day, when possible. Cross-fostering occurred twice (OVX-Prem, Sham-Prem), and sex-skewed litters were maintained twice (Sham-Oil, OVX-Prem). Rats were left undisturbed except for cage changing and weighing which occurred weekly during gestation. Two rats assigned to the primiparous group failed to become pregnant (one Sham-Prem, one Sham-Oil); one animal failed to lactate (Sham-Prem); and one animal cannibalized her litter (OVX-Oil). These rats were excluded from the study.

**Figure 1.**
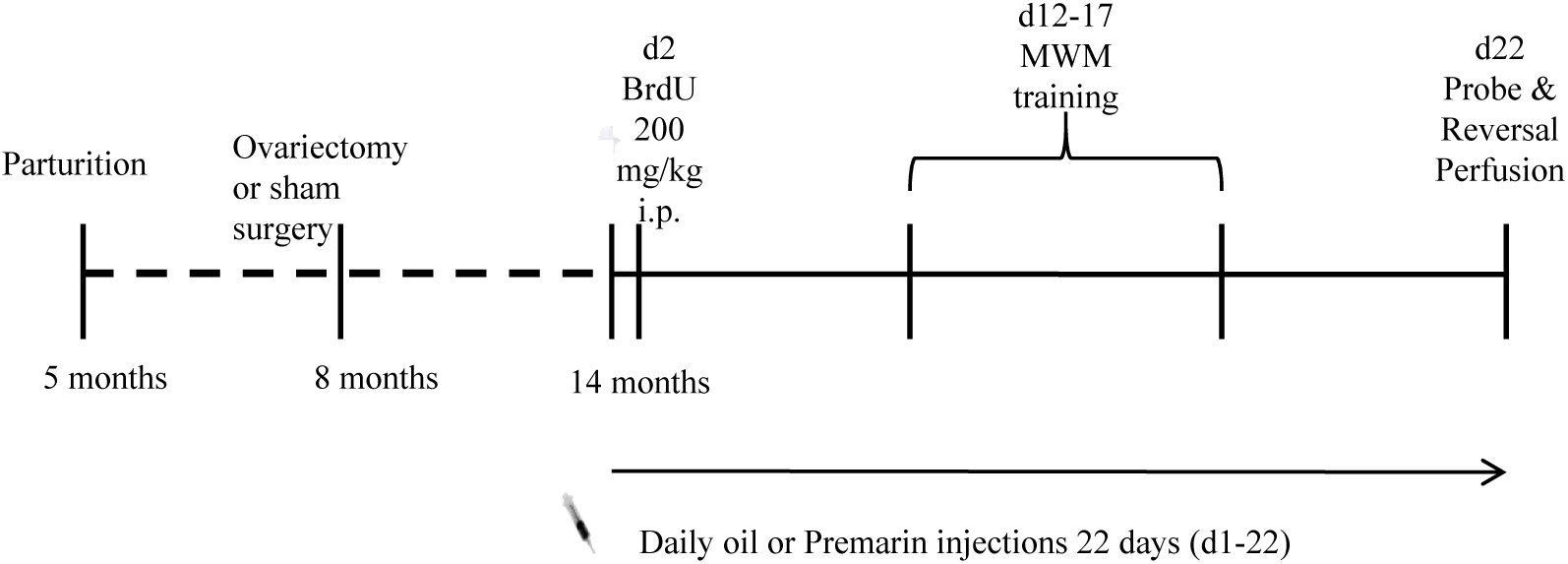
Experimental outline. MWM = Morris Water Maze, BrdU= bromodeoxyuridine.

### Maternal behavior observations

Observations of maternal behaviors were done three times per day from postpartum days 2 through 8. Observations were taken between 9:00-10:30am, 1:00-2:30pm, and 5:00-6:30pm daily, during which each dam was observed for 10 continuous minutes. The amount of time spent engaging in the following behaviors was recorded: licking and grooming, nursing (arched-back, blanket, passive), and off nest behaviors (including self-grooming and sleeping).

### Ovariectomy and Sham Surgery

At 8 months of age, rats underwent bilateral ovariectomy or sham surgery. Rats were induced and anesthetized with isofluorane, which was delivered at an induction flow rate of 5% in 1.5% oxygen. Rats were maintained at a surgical plane of anesthesia on a warming blanket using an isofluorane flow rate of 1.5-2.5% and were given an injection of Lactated Ringer’s Solution (10 mL) to maintain fluid balance, and Ketoprofen (0.5 mL/kg; Anafen, MERIAL Canada Inc., Baie d’Urfe’, Quebec, Canada) and Marcaine (0.10 mL; Bupivacaine, HOSPIRA Inc., Lake Forest, Illinois, Canada) as analgesics. Sham surgery consisted of skin and muscle incisions that were subsequently sutured without damage to or manipulation of the ovaries. Rats were single-housed for 7 days or until their sutures completely healed, at which point they were re-paired with cage mates.

Between the time of parturition and behavioral testing (from 5 months of age to approximately 14 months of age), 17 animals reached humane endpoint due to various factors common with aging (e.g., respiratory distress, malocclusion, urogenital or mammary tumor), or developed ulcerated foot sores that precluded participation in behavioral testing (Null-Sham-Oil: 2; Null-Sham-Prem: 2; Null-OVX-Oil: 2; Null-OVX-Prem: 1; Prim-Sham-Oil: 4; Prim-OVX-Oil: 1; Prim-Sham-Prem: 3; Prim-OVX-Prem: 2).

### Hormone Administration

At 14-15 months of age, rats began a daily subcutaneous injection of either Premarin (20μg Conjugated equine estrogens (CEE)/0.1 ml sesame oil; Wyeth Pharmaceuticals, Markham, ON, Canada) or sesame oil (equivalent volume) and continued for 22 days. Injections were given between 7:30 and 9:30 am, at least 2 hours before any behavioral testing. The dose of Premarin was based on Acosta et al (2009) to correspond to the most common daily dose taken by women (0.625mg; .00893 mg drug/kg body weight for a woman of average weight). The equivalent injected dose in rats has previously been found to influence memory (Acosta et al., 2009).

### 5-bromo-2-deoxyuridine (BrdU) Administration

On the second day of hormone injections, rats received a single i.p. injection of 200mg/kg 5-bromo-2-deoxyuridine (BrdU: Sigma, St. Louis, MO). The BrdU solution was prepared to a concentration of 20mg/ml just prior to injection by dissolving BrdU in freshly prepared warm (< 40 °C) 0.9% saline containing 0.7% 1N NaOH. BrdU is a thymidine analogue that incorporates into the DNA of dividing cells during the synthesis phase of the cell cycle within two hours after administration (Packard, et al., 1973). Depending on the amount of time elapsed between injection and perfusion of the animal, BrdU can be used to assess cell proliferation or cell survival in the dentate gyrus (Taupin, 2007). In the current study, rats were perfused 21 days after BrdU administration and 22 days after first hormone or oil injection. Therefore, the number of BrdU-labeled cells measures the survival of adult-generated hippocampal cells over a 21 day period after being produced and maintained under Premarin or oil administration.

### Lavage

Phase of the estrous cycle influences both hippocampal plasticity (Tanapat et al., 1999; Wooley et al., 1990) and spatial learning (Warren & Juraska, 1997). Furthermore, the length, and regularity of the estrous cycle changes with aging in rats (LeFevre & McClintock, 1988). Therefore, estrous cycles were monitored in our study by vaginal lavage. Lavage samples were transferred onto microscope slides, stained with Cresyl Violet, and left to dry. A rat was determined to be in the proestrous stage if at least 70% of cells were nucleated epithelial cells (Byers et al., 2012). Vaginal lavage samples were qualitatively categorized for evidence of normal cycling, abnormal cycling (consecutive lavage-cycles varying in length or order), persistent diestrus (consecutive lavage-cycles consisting primarily of leukocyte-dense cell samples), or persistent estrus (consecutive lavage-cycles consisting primarily of cornified cells; modified from Levefre & McClintock, 1988 to reflect the shorter sampling period in this study). Of the 32 sham rats that completed the study, 11 had irregular cycling, 11 were in persistent estrus and 10 were cycling normally (See Table 1).

**Table 1.** Percentage of sham-operated rats exhibiting regular cycling, or irregular cycling. A chi-square on regular vs irregular cycling was significant χ^2^=7.89, p=0.048.

|  | % regular cycling | % irregular cycling |
| --- | --- | --- |
| Nulliparous - oil | 66.67(6) | 33.33(3) |
| Nulliparous - Premarin | 12.50 (1) | 87.50 (7) |
| Primiparous - oil | 12.50 (1) | 87.50 (7) |
| Primiparous - Premarin | 28.57 (2) | 71.43 (5) |

### Morris Water Maze Training

Morris Water Maze training was conducted on injection days 12-17 during 9am −2 pm each day, with animals counterbalanced. There were four trials each day of 60s in duration or until the rat located the hidden platform. If the rat did not find the platform within 60s, the experimenter guided it to the platform, where it stayed for 10s. The inter-trial interval was approximately 7 minutes. Within each session, trials were randomly started from four different points within the pool, which were never repeated within a training day. The first day of water maze training was a visible platform test. A visible platform (white, 2 cm above the water) was placed in the SE quadrant of the maze. Days 2-6 of training used the standard reference memory version of the water maze; the platform was submerged 2 cm below the pool surface and remained in the NE quadrant throughout training. Total distance, swim speed and latency to reach the hidden platform were calculated for each day across the four trials using ANY-maze software.

### Probe Trial, Reversal Learning, and Perfusion

Five days after the last training trial and 21 days after BrdU injection, rats received a 60-second probe trial, during which the platform was removed from the pool. Percentage of time spent in the platform zone was recorded. The platform zone was defined as a circular region centered on the hidden platform encompassing 5% of the area of the water maze. Approximately 10 minutes after the probe trial, rats underwent 4 reversal trials in which the hidden platform was moved to the SW quadrant. Ninety minutes after the probe trial, rats were weighed and administered an overdose of sodium pentobarbitol and were perfused transcardially with 60 ml 0.9% saline followed by 120 ml 4% paraformaldehyde (Sigma-Aldrich). Just prior to perfusion, ovaries (sham rats only), uterine and adrenals were extracted and weighed (organ mass was divided by body mass to calculate a ratio and used in analyses). Due to an error, fewer animals had adrenals collected (n=4-6 per group). Brains were extracted and post-fixed in 4% paraformaldehyde overnight, transferred to 30% sucrose (Fisher Scientific) solution 24h later, and remained in sucrose solution at 4°C until sectioning. Brains were sliced using a Leica SM2000R microtome (Richmond Hill, Ontario, Canada) into 4μm coronal sections. Sections were collected in series of every 10th section throughout the entire rostral-caudal extent of the hippocampus and stored at −20°C in a cryoprotective solution consisting of 30% ethylene glycol (Sigma-Aldrich, St. Louis, MO, USA) and 20% glycerol (Sigma-Aldrich) in 0.1 M phosphatebuffer (PB, pH 7.4).

### Hormone Assays

Blood was taken at the time of perfusion from the right atrium. Blood samples were stored overnight at 4°C and centrifuged at 10,000*g* for 15 minutes. Serum was collected and stored at - 20°C. To determine the concentration of estradiol and estrone in circulation, radioimmunoassay (RIAs) were conducted in duplicate on serum according to manufacturer’s instructions (Beckman Coulter, Mississauga, Ontario, Canada). The sensitivity for the ultrasensitive 17β-estradiol kit is 2.2 pg/mL and the antibody is highly specific for 17β-estradiol, with 2.40 % cross-reactivity with estrone. The sensitivity for the estrone kit is 1.2pg/mL and has 1.25 % cross-reactivity with 17β-estradiol. Average intra-assay coefficients of variation were 10.66% and 10.73% for the estrone and 17β-estradiol kit, respectively.

### Serum cytokine quantification

Serum cytokines were quantified using a multiplex electrochemiluminescence immunoassay kit (V-PLEX Proinflammatory Panel 2, Rat) from Meso-Scale Discovery (Rockville, MD), used according to manufacturer instructions. Samples were run in duplicates and the following cytokines were quantified simultaneously in each sample: Interferon gamma (IFN-γ), Interleukin-1beta (IL-1β), Interleukin-4 (IL-4), Interleukin-5 (IL-5), Interleukin-6 (IL-6), KC/GRO, Interleukin-10 (IL-10), Interleukin-13 (IL-13), and tumor necrosis factor alpha (TNF-α). A Sector Imager 2400 (Meso Scale Discovery) was used for plate reading, and the Discovery Workbench 4.0 software (Meso Scale Discovery) was used for data analyses. Lower limits of detection (LLODs) were as follows (pg/ml): IFN-□: 0.163; IL-1β: 1.48; IL-4: 0.179; IL-5: 7.64; IL-6: 2.4; IL-10: 0.233; IL-13: 0.78; TNF-α: 0.156; and KC/GRO: 0.085.

### Immunohistochemistry

#### DCX

One series of brain tissue was labeled for the immature neuronal marker doublecortin (DCX). Tissue was pretreated with 0.6% hydrogen peroxide for 30 minutes at room temperature after rinsing in 0.1 M PBS. It was then incubated at 4°C for 24 h in a primary antibody solution consisting of 1:1000 polyclonal goat anti-doublecortin (Santa Cruz Biotechnology, Santa Cruz, CA, USA, Cat# sc-8066, RRID:AB_2088494), 0.04% Triton-X, and 3% normal rabbit serum, dissolved in 0.1 M PBS. The tissue was washed in 0.1 M PBS and then incubated for 24 h at 4°C in a secondary antibody solution containing 1:500 biotinylated rabbit anti-goat (Vector Laboratories, Burlingame, CA, USA) in 0.1 M PBS. Tissue was then incubated for 4 h at room temperature in an avidin-biotin solution containing 1:1000 avidin and 1:1000 biotin in 0.1M PBS (ABC kit; Vector Laboratories), rinsed in PBS, and washed in 0.175 M sodium acetate buffer. Doublecortin-expressing cells were visualized by developing tissue for approximately 10 minutes in a diaminobenzidine (DAB; Sigma Aldrich) solution. Once staining was complete, tissue was mounted on glass slides, dehydrated, cleared with xylene, and coverslipped using Permount (Fisher Scientific).

#### Zif268

A series of hippocampal sections was stained for zif268, an immediate early gene. Tissue was rinsed in 0.1M PBS overnight and 3 x10 minutes the following day. The tissue was incubated in 0.6% H_2_O_2_ for 30 minutes at room temperature and re-rinsed. The tissue was then incubated in the primary antibody, 5abbit anti-Erg-1 ((Santa Cruz Biotechnology Cat# sc-189, RRID:AB_2231020)) at 4°C for 18 h with 0.04% Triton-X and 3% normal goat serum (NGS; Vector Laboratories) in 0.1M PBS. Next, the tissue was rinsed in PBS and incubated in the second antibody solution (goat anti-rabbit Biotinylated IgG in 0.1M PBS; 1:1000; Vector Laboratories, Burlington, ON, Canada) for 18 h at 4°C. Tissue was then rinsed in 0.1M PBS and incubated for 1 h at room temperature in an avidin-biotin complex dissolved in 0.1M PBS, as per instructions in the ABC kit (Vector Laboratories). The tissue was rinsed first in 0.1M PBS followed by a rinse in sodium acetate. Brain sections were then transferred to a DAB solution and incubated for 5 minutes in a dark room, then rinsed with sodium acetate followed by 0.1M PBS. The tissue was mounted onto glass microscope slides, dried overnight, dehydrated in ethanol, cleared with Xylene, and cover-slipped with Permount.

#### BrdU/zif268

Immunofluorescent double-labelling of BrdU and zif268 was conducted to quantify the percentage of 21-day old neurons activated during spatial memory. Staining began with three rinses for 10 minutes each in 0.1M PBS. The tissue was then incubated for 24 h in the zif268 primary antibody solution containing rabbit anti-zif268 (1:1000; Egr-1 SC-189, Santa Cruz Biotechnology, Santa Cruz, CA, USA), with 4% normal donkey serum, 0.03% Triton X, and 0.1M PBS. The slices were then rinsed three times for 10 minutes in 0.1M PBS. Tissue was then incubated in the secondary antibody donkey anti-rabbit Alexa 488 (1:500; Invitrogen Molecular Probes, Oregon, USA) for 18 h at 4°C. The tissue was then fixed in 4% paraformaldehyde for 10 minutes, washed twice in 0.9%NaCl for 10 minutes each followed by 2N HCl at 37°C for 30 minutes. Next tissue was rinsed three times in 0.1M PBS and incubated in mouse anti-BrdU (1:500; Roche Diagnostics GmbH, Mannheim, Germany, Roche Cat# 11170376001, RRID:AB_514483) with 4% normal donkey serum, and 0.03% Triton X in PBS for 24 h at 4°C. The tissue was then rinsed three times for 10 minutes in 0.1M PBS after which it was incubated in donkey anti-mouse Cy3 (1:250: Jackson Immuno Research Laboratories Inc., Philadelphia, USA) for 18 h. To conclude, the sections were rinsed three times for five minutes each, mounted on glass slides, and coverslipped using PVA DABCO.

#### BrdU/NeuN

A final series of hippocampal sections was double labelled for BrdU and NeuN, a mature neuronal protein, to quantify the proportion of adult-generated hippocampal cells with neuronal morphology. Sections were incubated in 0.1M TBS containing the primary antibody, 1:250 mouse anti-NeuN (EMD Millipore, Cat# MAB377, RRID:AB_2298772) and BrdU (BioRad / AbD Serotec Cat# OBT0030S, RRID:AB_609570) at 4°C for 48 h. The tissue was rinsed for 10 minutes three times in TBS. Tissue was then incubated in the secondary antibody, donkey anti-mouse Alexa 488 (1:200; Invitrogen Molecular Probe, Burlington, ON, Canada), for 18 h. The tissue was then fixed using 4% paraformaldehyde for 10 minutes, rinsed two times for 10 minutes in 0.9% NaCl, and was incubated in 2N HCl at 37°C for 30 minutes. Tissue was then incubated in rat anti-BrdU (1:500; AbD Serotec, Raleigh, NC, USA) for 48 h at 4°C, and then incubated for 24 h in donkey anti-rat Cy3 (1:500; Jackson ImmunoResearch; PA, USA). The tissue was then rinsed three times for 10 minutes each in TBS, mounted on glass slides, and cover slipped using PVA DABCO.

### Cell counting

All cell counting was performed by an experimenter blind to treatment groups. DCX-expressing cells were counted in every 10^th^ section of the hippocampus using a Nikon E600 light microscope under a 40x objective lens as described elsewhere (Barha and Galea, 2013, Barha et al., 2011b, Brummelte and Galea, 2010 and Epp et al., 2011). BrdU labelled cells were counted in every 10^th^ section using an Olympus CX22 microscope with a 100x objective. Cells were considered to be BrdU-labeled if they were intensely stained and exhibited medium-sized round or oval cell body (Cameron, et al., 1993). The percentages of BrdU labeled cells co-expressing zif268 or NeuN were calculated by identifying, exhaustively, whether BrdU-labelled cells were co-labeled with zif268 or NeuN. Area measurements for the GCL were obtained with digitized images and the software ImageJ (NIH).

Volume estimates of the dentate gyrus were calculated using Cavalieri’s principle (Gundersen & Jensen, 1987). Total number and density of BrdU-labeled and DCX-expressing cells in each region (Dorsal or Ventral) were calculated. We noted whether immunoreactive cells were located in the dorsal or ventral GCL using the criterion defined by Banasr et al. (2006), as spatial memory is more strongly associated with the dorsal hippocampus and anxiety/stress behavior with the ventral hippocampus (See Fanselow & Dong, 2010 for review). DCX-expressing cells were classified into stages of maturity based on Plümpe et al. (2006). Type 1 cells (proliferative stage cells) included DCX-expressing cells with no processes or short, plump processes no longer than a cell width; Type 2 cells (intermediate stage cells) had unbranched processes of intermediate length reaching no farther than the moleculayer; Type 3 cells (postmitotic stage cells) had a more mature appearance, with denritic branching in the molecular and/or granule cell layer.

### Zif268 Optical Density

For each animal, eight sections of the hippocampus (four ventral, four dorsal) were examined for zif268 expression. Images of these sections were taken at 40X magnification using a Nikon E600. These images were converted to 8-bit images (grey scale) and the optical density of zif268-expressing cells was quantified using Image J (Rasband, W.S., ImageJ, U.S. National Institutes of Health, Bethseda, Maryland, USA). From each segment, an ellipse of 50 μm area was sampled from the dentate gyrus, the CA1 hippocampal field, and the CA3 hippocampal field. To accommodate differences in staining between tissue samples, staining intensity was assessed by quantifying Zif268 staining in the tissue background (using the corpus callosum). Therefore, a fourth ellipse was sampled from the background of the image. The optical density (OD) of each region of interest was expressed as the mean gray value per μm^3^. The background measure was subtracted from the OD in the regions of interest (DG, CA1, CA3) to obtain an OD measure that compensates for differences in intensity of staining.

### Data analyses

All analyses were conducted using Statistica (Statsoft Tulsa, OK). Analysis of variance (ANOVA) were used on most dependent variables of interest with ovarian status (OVX, sham), hormone treatment (Premarin, oil) and parity (nulliparous, primiparous) as between-subjects variables. For some analyses, training day (1 to 5), DCX cell maturity stage, region (dorsal, ventral), or hippocampal field (CA1, CA3, DG) were used as the within-subjects variables. Post-hoc tests used Newman-Keuls. Any a priori comparisons were subjected to a Bonferroni correction. Pearson product-moment correlations were performed on dependent variables of interest. The significance level was set at α=0.05

## Results

### Among primiparous rats, surgery and treatment groups were similar in their previous maternal behavior towards offspring

Treatment and ovarian status groups did not differ significantly in maternal behavior (See Table 2). There were no significant main effects or interactions between ovarian status or treatment on active (nursing, licking) versus passive/off-nest behaviors recorded on post-natal days 2-8 (*p*’s > 0.11).

**Table 2.** Mean(± SEM) percentage of time spent in active maternal behaviors versus passive or off-nest behaviors during postnatal days 2-8.

|  | Nursing/Licking | Passive/Off-Nest |
| --- | --- | --- |
| Sham - oil | 45.31 ± 5.64 | 54.69 ± 5.64 |
| Sham - Premarin | 58.33 ± 3.31 | 41.67 ± 3.31 |
| OVX - oil | 53.16 ± 4.97 | 46.84 ± 4.97 |
| OVX - Premarin | 51.45 ± 3.40 | 48.55 ± 3.40 |

### Distance to reach the visible platform did not differ across groups

Two rats (Null-Sham-Oil and Null-Sham-Prem) were unable to find the visible platform during this phase of testing and were excluded from further testing. Rats improved significantly across visible platform training trials (main effect of trial, *F*(3, 204) = 6.922, *p* < 0.001; data not shown), but there were no other significant main or interaction effects (p’s > 0.31).

### Premarin treatment impaired spatial reference memory performance in primiparous rats and enhanced spatial reference memory performance in nulliparous rats

As expected, distance travelled during reference memory training decreased significantly across days, *F*(4, 272) = 71.39, *p* < 0.0001 (data not shown). There was a significant parity by treatment interaction in distance to reach the hidden platform, F(1, 68) = 4.16, *p* = 0.045. However, post-hoc tests did not reveal any significant differences between groups (all *p*’s > 0.19). Latency to reach hidden platform decreased across days, F(4, 272) = 77.47, *p* < 0.0001, but there were no group differences in latency to reach the hidden platform (*p*’s > 0.13). Groups did not differ in swim speed across days (*p*’s > 0. 19).

Other researchers have found effects of hormone administration and/or parity differ between early and late stages or blocks of learning (Frye, 1995; Barha et al., 2015; Barha & Galea, 2013; Acosta et al., 2009; 2010). To examine whether there were significant differences in total distance travelled during early versus late learning, we summed the total distance across trials for days 1-3 and 4-5. Premarin-treated primiparous rats had longer distances to reach the platform than Premarin-treated nulliparous rats (*p* = 0.032; parity by treatment F(1,68) = 5.655, *p* = 0.020). *A priori* we thought there would be differences in parity and treatment groups across days, and examining total distance from days 1-3 versus days 4-5, we found that Premarin-treated primiparous rats travelled longer distances to reach the platform than Premarin-treated nulliparous rats across days 1-3 but not 4-5 (*p*’s = 0.018 and 0.46, respectively; Figure 2A and B). Furthermore, Premarin treatment increased distances to reach the hidden platform on days 1-3 compared to oil treatment in primiparous rats (*p* = 0.021) but decreased distances to reach the hidden platform in nulliparous rats (*p* = 0.027; see Figure 2A and 2B).

**Figure 2.**
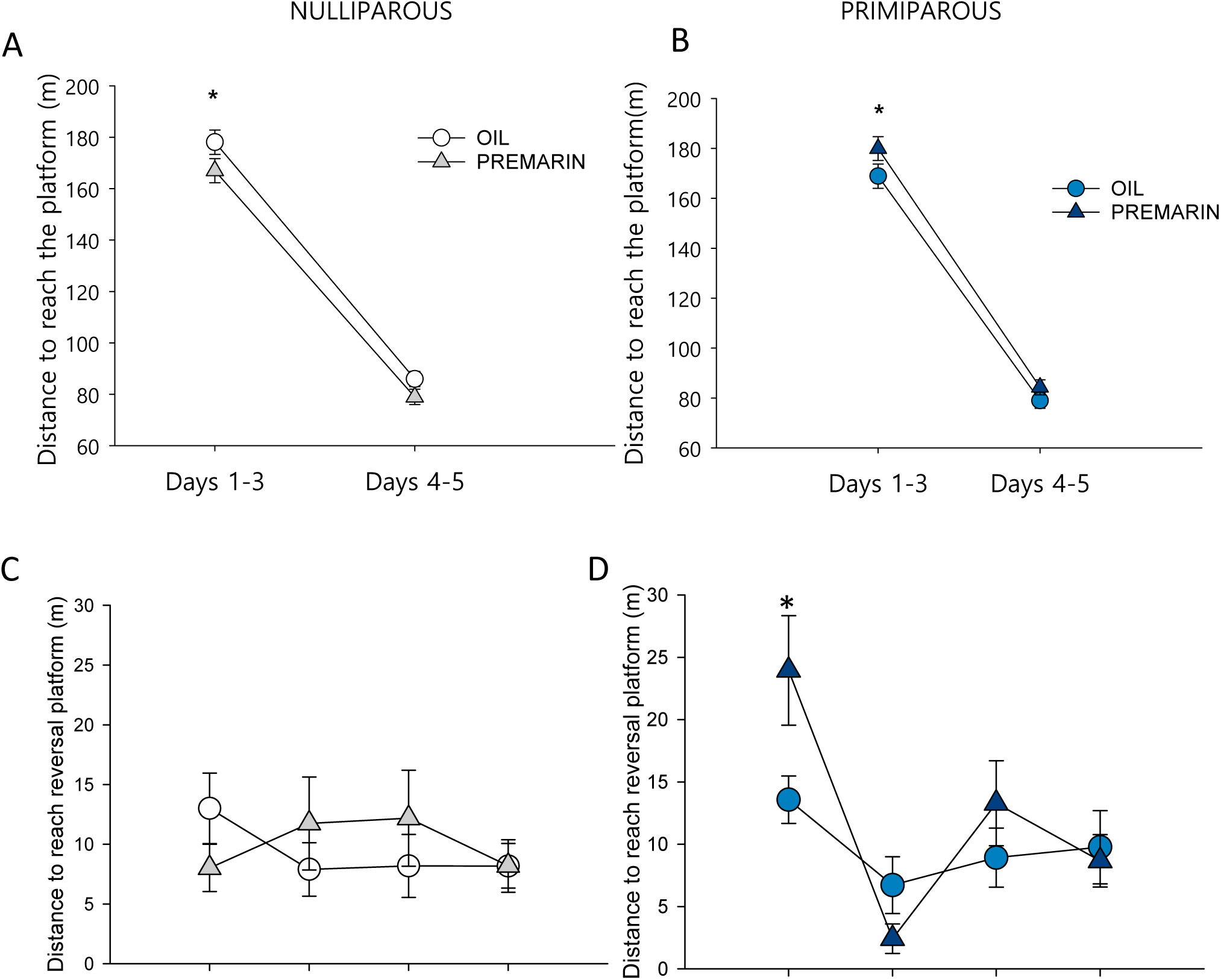
Spatial learning as measured by distance to reach the hidden platform (m) in the Morris Water Maze. Premarin HT reduced distance to reach the hidden platform in nulliparous rats (A) but increased distance to reach the hidden platform in primiparous rats (B) <u>Reversal Learning</u>: Under sham surgery, Premarin treatment increased distance to reach the hidden platform on the first trial compared to oil in Primiparous rats (D) but there were no significant differences with Premarin in nulliparous rats (C). Under ovariectomy, there were no group differences in distance to reach the hidden platform over trials (E).

### Under Premarin treatment, sham-operated primiparous rats have impaired performance on the first reversal trial compared to sham-operated nulliparous rats

Under Premarin treatment, primiparous sham-operated rats had greater distance to find the new platform location on the first reversal trial compared to all other groups (*p*’s < 0.007; parity by ovarian hormone status by treatment by trial interaction: *F*(3,201) = 3.93, *p* = 0.009; See Figure 2D). There were also significant main effects of trials [*F*(3,201) = 27.19, *p* < 0.001], surgery [*F*(1,67) = 5.62, *p* = 0.021] and interactions (trial by parity and trial by ovarian hormone status by parity, *F*(3,201) > 119.46, *p*’s < 0.039). Under ovariectomy, there were no group differences in distance to reach the hidden platform over trials.

### Premarin improved spatial memory retrieval in sham-operated nulliparous rats. Ovariectomy improved spatial memory retrieval in primiparous rats

We expected *a priori* that reproductive experience would alter the effects of Premarin and perhaps ovarian hormone status on spatial memory retrieval. *A priori* analyses indicated that under sham surgery, nulliparous but not primiparous had improved probe trial performance with Premarin relative to oil (*p*’s = 0.05 and 0.37, respectively; Figure 3). Under ovariectomy, neither parity group was affected by Premarin (*p*’s > 0.44), but Premarin-treated primiparous females spent a greater percentage of time in the platform zone than Premarin-treated nulliparous rats *p* = 0.025; Figure 3). No other pairwise comparisons between parity groups or treatment groups were significant, *p*’s > 0.26. There were no other significant effects (all *p*’s ≥ 0. 19).

**Figure 3.**
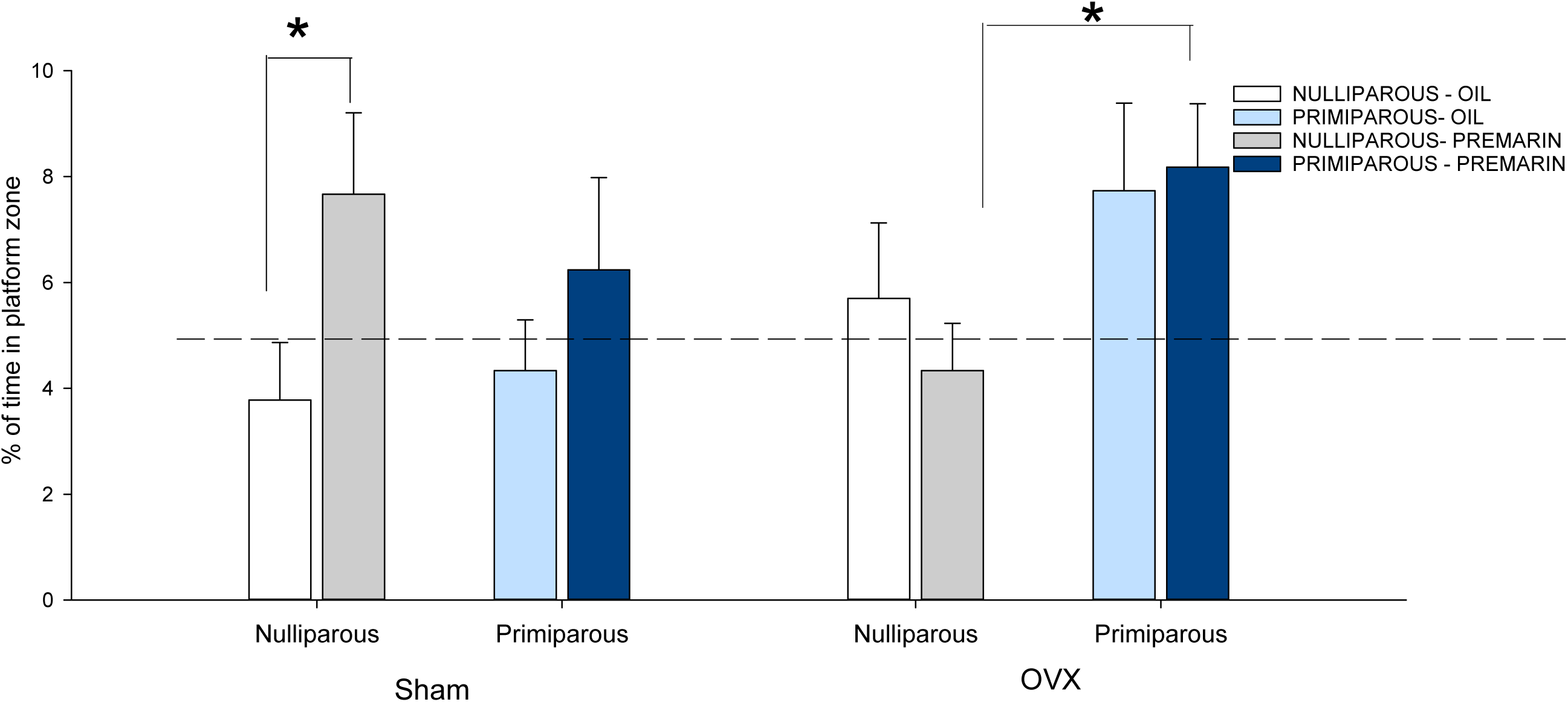
Time spent (%) in the platform zone during probe trial. Under sham surgery, nulliparous rats spent significantly more time in the platform zone under Premarin treatment compared to oil treatment. Under ovariectomy, Premarin-treated primiparous females spent significantly more time in the platform zone than Premarin-treated nulliparous rats. Dashed line indicates chance level. * denotes *p* < .05.

### Primiparous rats had greater number of BrdU-labelled cells than nulliparous rats. Premarin decreased the survival BrdU-labelled cells in the ventral dentate gyrus but increased it in the dorsal dentate gyrus in sham-operated females

There were no significant differences in volume for the dorsal or ventral dentate gyrus (all p’s >0.09; data not shown) and thus, estimated total counts are used. Primiparous rats had more BrdU-labelled cells than nulliparous rats (F (1,66)=4.712, p<0.03; see Figure 4A). In sham-operated females, Premarin increased the number of BrdU-labelled cells in the dorsal GCL (*p* = 0.02) but decreased the number of BrdU-labelled cells in the ventral GCL (Region by ovarian hormone status by Treatment interaction: *F*(1,66) = 19.34, *p* < 0.001), but there was no significant effect of Premarin in OVX groups; Figure 4B). Ovariectomized oil-treated rat had higher levels of BrdU in the dorsal dentate gyrus (p=0.004) but lower levels of BrdU-labeled cells in the ventral dentate gyrus (p=0.019) compared to Sham-oil treated females. The majority of BrdU-labelled cells co-expressed NeuN (>83%), indicating they were new neurons, and there were no significant differences as a function of parity, treatment, or surgery (all *p*’s > .20, see Table 3). The percentage of BrdU-labelled cells that co-expressing zif268 was not significantly different among groups (all other *p*’s ≥ 0.09; Table 3).

**Figure 4.**
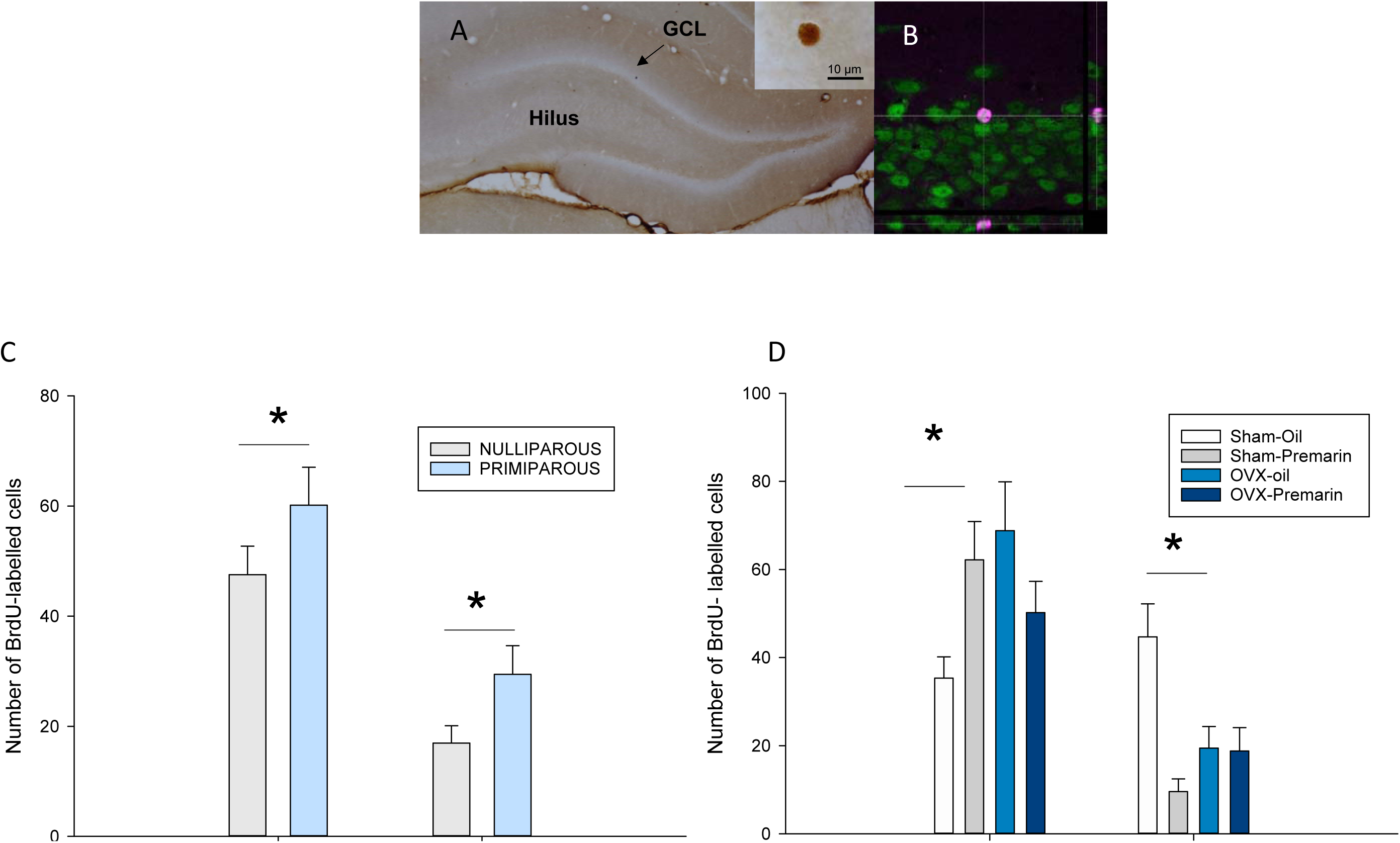
A. Primiparous rats had more BrdU-labelled cells than nulliparous rats surviving 21 days (+ SEM) in the dorsal and ventral GCL. **B.** Premarin treatment decreased the survival BrdU-labelled cells in the ventral dentate gyrus, but increased survival in the dorsal dentate gyrus in sham-operated females. **C.** Representative photomicrograph of a BrdU-labelled cell in a primiparous female, scale bar = 10μm. **D**. Confocal image of a BrdU/NeuN co-labelled cell, green=NeuN, red=BrdU. OVX: ovariectomy, BrdU=bromodeoxyuridine.

**Table 3.**
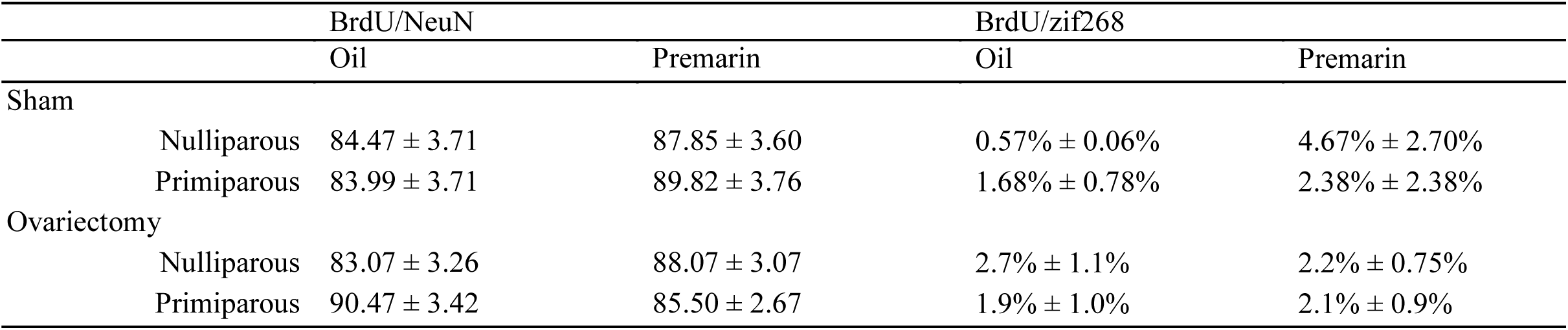
Mean(± SEM) percentage of BrdU-labelled cells co-expressing NeuN or zif268 in response to spatial memory retrieval.

|  |  | BrdU/NeuN |  | BrdU/zif268 |  |
| --- | --- | --- | --- | --- | --- |
|  |  | Oil | Premarin | Oil | Premarin |
| Sham |  |  |  |  |  |
| | Nulliparous | 84.47 ± 3.71 | 87.85 ± 3.60 | 0.57% ± 0.06% | 4.67% ± 2.70% |
| | Primiparous | 83.99 ± 3.71 | 89.82 ± 3.76 | 1.68% ± 0.78% | 2.38% ± 2.38% |
| Ovariectomy |  |  |  |  |  |
| | Nulliparous | 83.07 ± 3.26 | 88.07 ± 3.07 | 2.7% ± 1.1% | 2.2% ± 0.75% |
| | Primiparous | 90.47 ± 3.42 | 85.50 ± 2.67 | 1.9% ± 1.0% | 2.1% ± 0.9% |

### Primiparity increased DCX expression in the ventral dentate gyrus, and Premarin treatment reduced DCX expression dependent on region and parity

Premarin reduced DCX expression in the dorsal region in nulliparous rats (p=0.034) and in the ventral dentate gyrus in primiparous rats (*p* = .0007; region by parity by treatment interaction *F*(1,62) = 12.59, *p* = 0.0007; See Figure 5A). Primiparous oil-treated rats had higher expression of DCX in the ventral region than nulliparous oil-treated rats (p=0.014). There were also main effects of treatment and region (both *p*’s <0.034) but no other significant effects on DCX-expressing cells (p > 0.06).

**Figure 5.**
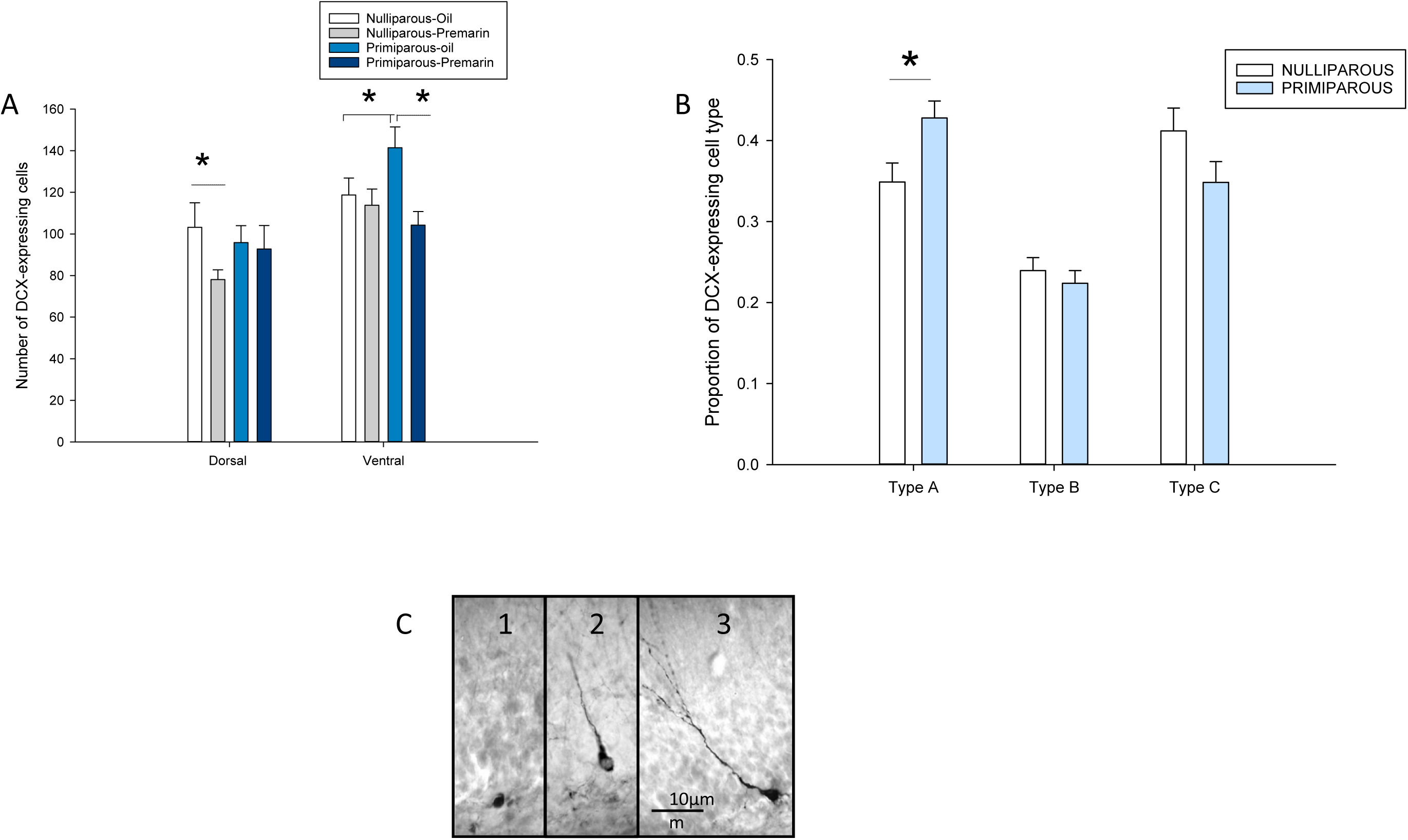
A. Premarin treatment decreased the density of doublecortin (DCX)-expressing cells in the ventral dentate gyrus of primiparous rats and in the dorsal dentate gyrus of nulliparous rats compared to oil-controls. B. Primiparity increased the proportion of Type 1 DCX-expressing cells compared to nulliparous rats (*p* = 0.044) but parity did not significantly affect the proportion of Type 3 cells (*p* = 0.10) or Type 2 cells (*p* = 0.69). C. Photomicrographs of Type 1, 2 and 3 DCX-expressing cells.

### Primiparity increased the proportion of proliferative immature neurons

Primiparity increased the proportion of Type 1 (proliferative) DCX-expressing cells compared to nulliparous rats (*p* = 0.044) but parity did not significantly affect the proportion of Type 3 (*p* = 0.10) or Type 2 cells (p = .67; See Figure 5B; stage by parity interaction, *F*(2,108) = 3.11, *p* = 0.048). There were no other significant main or interaction effects on DCX morphology (*p*’s > 0.07).

### Primiparous rats have increased zif268 expression in the CA3 region in response to spatial memory retrieval compared to nulliparous rats. Premarin increased ventral CA3 zif268 expression in primiparous but not nulliparous rats

Primiparous rats had greater zif268 expression in the CA3 region than nulliparous rats (main effect of parity (F(1, 57)= 4.13, p=0.047). Ovariectomy reduced zif268 expression in the ventral CA3 region, regardless of parity (F (1, 57) =7.40, p=0.009). A priori we expected differences with treatment and parity, and Premarin increased zif268 expression in the ventral CA3 in primiparous rats (p<0.00007) but not in nulliparous rats (p=0.811; see Figure 6).

**Figure 6.**
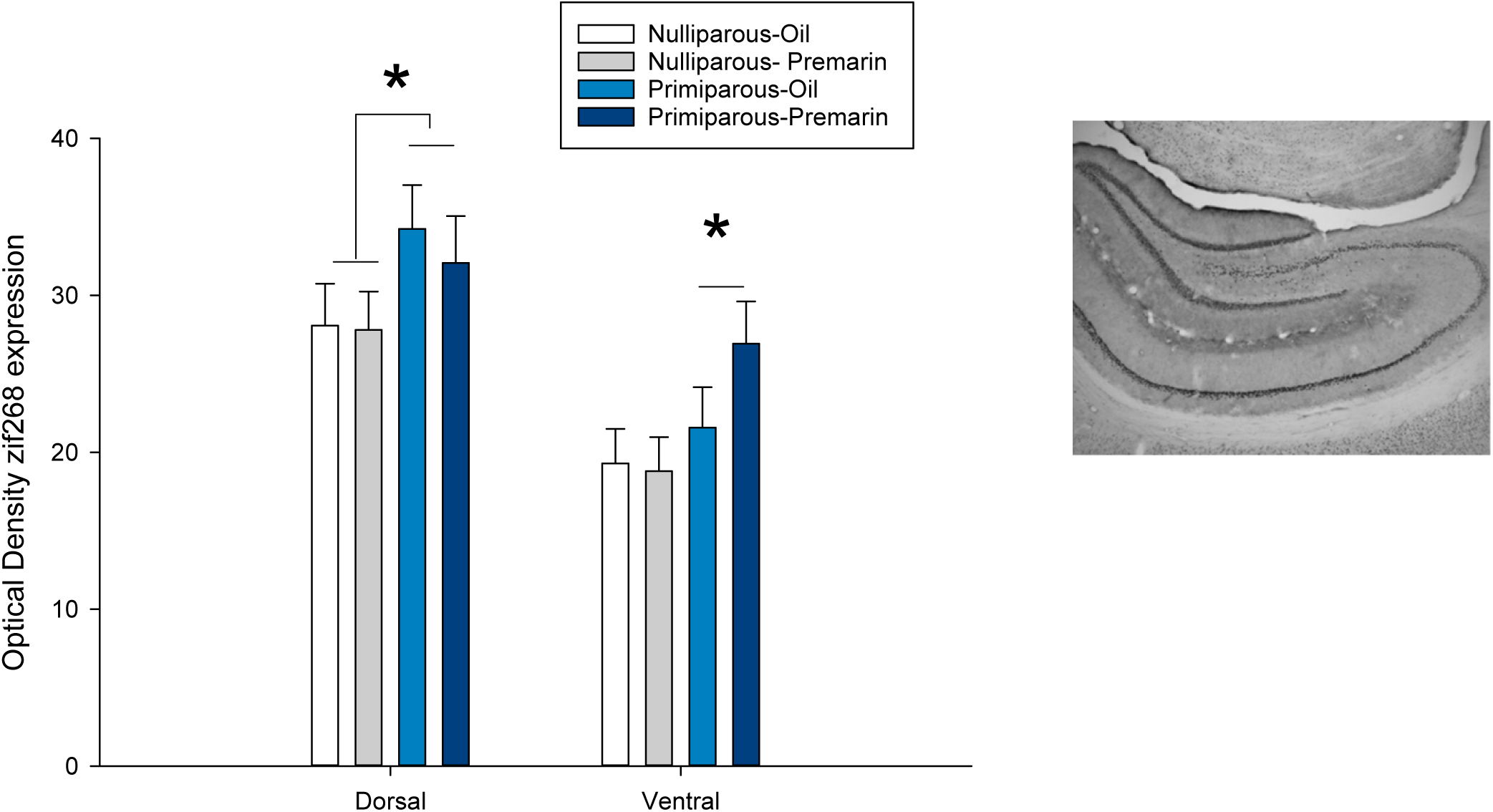
A Primiparity increased zif268 expression in the CA3 region in response to spatial memory, regardless of treatment, ovarian hormone status or region (main effect of parity). Furthermore, Premarin increased zif268 expression in the ventral CA3 region in response to spatial memory in primiparous but not nulliparous rats. * denotes *p* < .05.

In the CA1 region, there was greater zif268 expression in the dorsal versus ventral region, regardless of parity or treatment (main effect of region: F (1,57)=6.72, p=0.012). Again, ovarian hormone status reduced zif268 expression in the ventral CA1 region, regardless of parity or treatment (F(1, 57)=3.84, p=0.054; data not shown). There were no other significant differences in the CA1 region between groups (all p’s >0.17). There were no significant main or interaction effects on zif268 expression in the dentate gyrus (all p’s > 0.12; data not shown).

### Premarin increased TNF-α and KC/GRO in sham nulliparous, but not primiparous, rats

Premarin increased serum levels of TNF-α and KC/GRO in sham nulliparous rats compared to oil-treated nulliparous rats (p=0.006, p=0.048, respectively; see Figure 7). Premarin did not increase KC/GRO serum levels in primiparous rats (*p*’s >0.7) but there was a trend for Premarin to decrease TNFα levels in primiparous rats (p=0.0502). For TNF-α there was a significant interaction of parity by treatment by surgery (p= 0.024) while for KC/GRO this interaction was a trend only (p=0.07). Sham rats (250.0 ± 35.6) had higher serum levels of KC/GRO than OVX (167.7 ±18.5) rats (main effect of surgery: F(1, 32)=4.59, p=0.04). There were no other significant effects for TNF-α or KC/GRO. However, there was a trend for primiparous rats (8.34±1.87 pg/ml) to have higher serum levels of IL-6 compared to nulliparous rats (4.5±0.68 pg/ml; p=0.08). There were no other significant effects of surgery, Premarin or parity on any of the other cytokines: IL5, IL4, Ilβ, IL13, IL10, IFNγ (all p’s >0.16).

**Figure 7.**
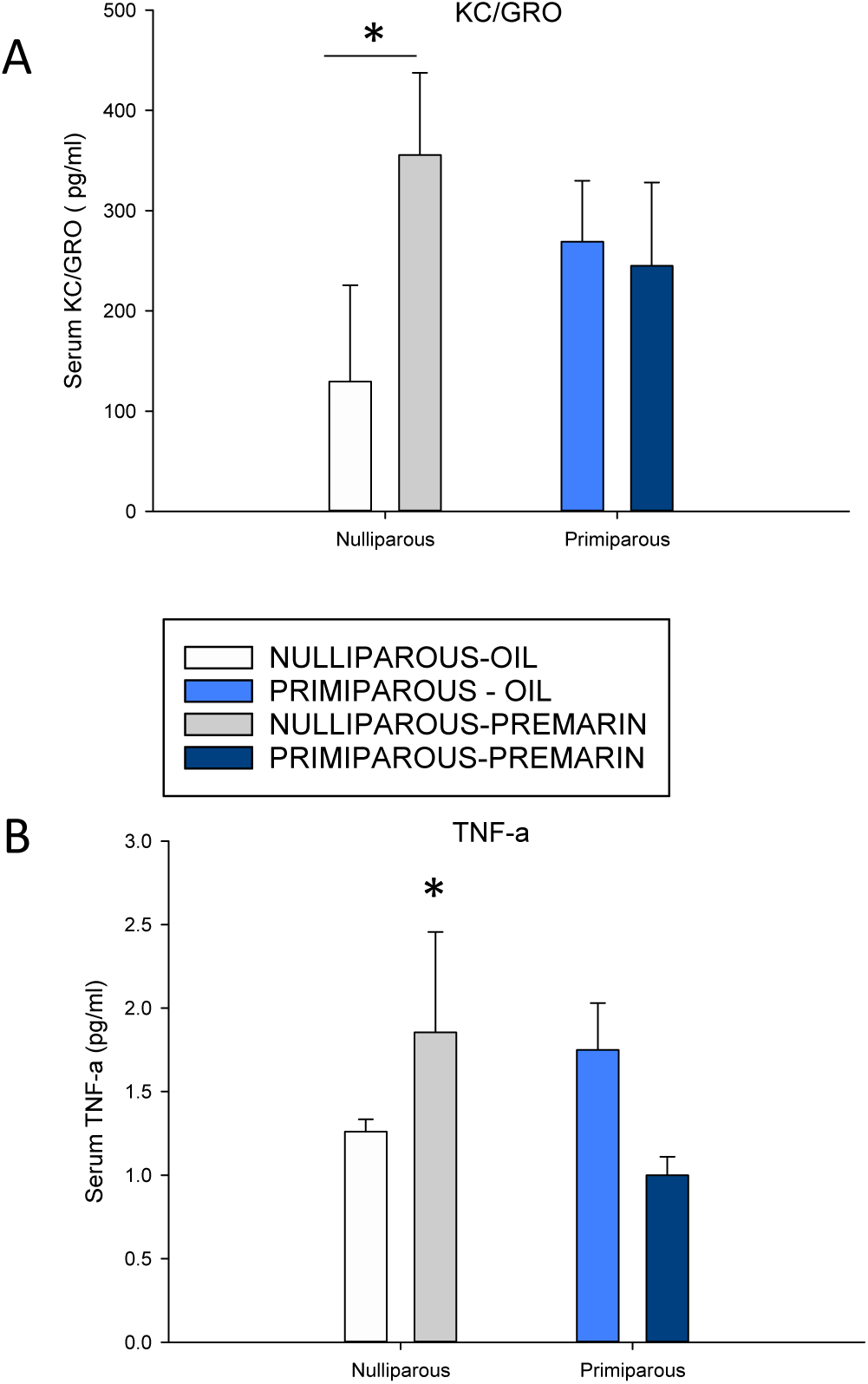
Premarin increased serum KC/GRO (A) and TNF-α (B) levels in sham nulliparous but not primiparous rats. * denotes *p* < .05 from oil-treated groups.

### Premarin prevents ovariectomy-induced decrease in relative adrenal mass in primiparous rats only. Premarin treatment reduced relative ovary mass regardless of parity

Ovariectomy significantly reduced relative adrenal mass (main effect of surgery: *F*(1,31) = 37.82, *p* < 0.0001). However a priori results indicated that ovariectomy reduced relative adrenal mass in all groups (*p*’s < 0.002) except in Premarin-treated primiparous rats (*p* = 0.47; parity by surgery by treatment interaction, *F* (1,31) = 3.88, *p* = 0.058; see Table 4).

**Table 4.** Mean(± SEM) organ mass to body mass ratio for adrenals, uterus, and ovaries

|  |  | Adrenals |  | Uterus |  | Ovaries |  |
| --- | --- | --- | --- | --- | --- | --- | --- |
|  |  | Sham | Ovariectomy | Sham | Ovariectomy | Sham | Ovariectomy |
| Oil |  |  |  |  |  |  |  |
| Nulliparous |  | 0.0165 | 0.0107 | 1.207 | 0.536 | 0.0215 | ----- |
| | | $\pm 0.001$ | $\pm 0.0004^*$ | $\pm 0.14$ | $\pm 0.15$ | $\pm 0.007$ | |
| Primiparous |  | 0.0170 | 0.0100 | 1.352 | 0.469 | 0.0216 | ----- |
| | | $\pm 0.001$ | $\pm 0.001^*$ | $\pm 0.23$ | $\pm 0.12$ | $\pm 0.006$ | |
| Premarin |  |  |  |  |  |  |  |
| Nulliparous |  | 0.0174 | 0.0108 | 0.920 | 0.620 | 0.0178 | ----- |
| | | $\pm 0.003$ | $\pm 0.001^*$ | $\pm 0.21$ | $\pm 0.06$ | $\pm 0.003$ | |
| Primiparous |  | 0.0147 | 0.0135 | 1.497 | 0.621 | 0.0194 | ----- |
| | | $\pm 0.002$ | $\pm 0.001$ | $\pm 0.22$ | $\pm 0.06$ | $\pm 0.005$ | |

\* Indicates significant difference from sham-operated rats

Premarin treatment reduced relative ovary mass compared to oil treatment (main effect of treatment: *F*(1,71) = 5.49, *p* = 0.022), regardless of parity (all interaction *p*’s > 0.50). As expected, ovariectomy significantly decreased relative uterine mass (main effect of surgery F(1,68) = 44.09, *p* < 0.0001). There were no other significant main or interaction effects (*p*’s > 0.06; see Table 4). As expected, ovariectomized rats were heavier than sham rats (main effect of surgery: *F*(1,68)=13.044, p<0.0006; data not shown), but there were no other significant effects on body mass at the time of perfusion.

### Premarin treatment increased estrone levels regardless of parity or surgery, and increased 17β-estradiol levels to a lesser degree. Primiparous rats had higher levels of estrone and were more likely to be irregularly cycling

As expected, Premarin increased both serum estradiol and estrone levels (treatment by estrogens interaction, *F*(1,67) = 38.96, *p* < 0.0001; both *p*’s < 0.0001) compared to oil treatment. Indeed, the increase in estrone levels was greater in magnitude than the increase with estradiol as the increase in estrone was 27x greater with Premarin while estradiol increased by 2-3x than control levels with Premarin. In addition, estrone levels were non-significantly greater than estradiol serum levels under oil treatment (*p* = 0.16) but were significantly greater than estradiol concentrations under Premarin treatment (*p* < 0.0001). There were no significant effects of parity or surgery on estrone or estradiol levels; however, this was likely due to the overwhelming treatment effects of Premarin. A separate analysis on oil-treated groups indicated that estrone levels were higher than estradiol levels (main effect of estrogens *F*(1,33) = 9.56, *p* = 0.004). A priori analyses revealed that parity groups did not differ in estradiol levels (*p* = 0.40) but did differ in estrone levels (*p* = 0.017), with primiparous females having greater estrone levels. Importantly there were no significant differences between estrone or estradiol levels between OVX and sham groups (*p* s >0.30; See Table 5). Nulliparous rats were more likely to be regularly cycling than primiparous rats (χ^2^=7.89, p=0.048; Table 1).

**Table 5.**
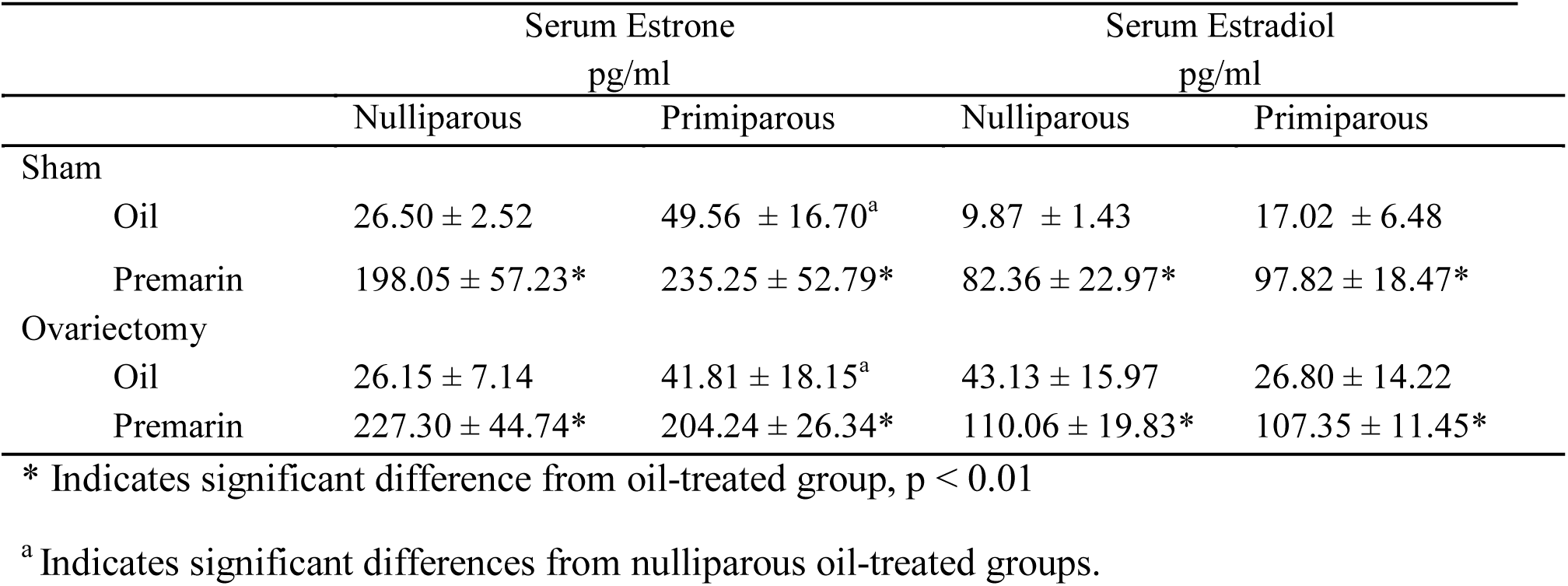
Mean(± SEM) serum estrone and estradiol levels (pg/ml) in sham-operated and ovariectomized nulliparous and primiparous rats receiving Premarin or oil treatment.

|  | Serum Estrone<br>pg/ml |  | Serum Estradiol<br>pg/ml |  |
| --- | --- | --- | --- | --- |
|  | Nulliparous | Primiparous | Nulliparous | Primiparous |
| Sham |  |  |  |  |
| Oil | 26.50 ± 2.52 | 49.56 ± 16.70<sup>a</sup> | 9.87 ± 1.43 | 17.02 ± 6.48 |
| Premarin | 198.05 ± 57.23* | 235.25 ± 52.79* | 82.36 ± 22.97* | 97.82 ± 18.47* |
| Ovariectomy |  |  |  |  |
| Oil | 26.15 ± 7.14 | 41.81 ± 18.15<sup>a</sup> | 43.13 ± 15.97 | 26.80 ± 14.22 |
| Premarin | 227.30 ± 44.74* | 204.24 ± 26.34* | 110.06 ± 19.83* | 107.35 ± 11.45* |

\* Indicates significant difference from oil-treated group, $p < 0.01$
<sup>a</sup> Indicates significant differences from nulliparous oil-treated groups.

### Serum cytokine levels (IL4, IL6, IL10 and IL13) were positively correlated with neurogenesis in the Premarin-treated Primiparous group only

Premarin was associated with positive correlations between cytokines and BrdU-labelled cells in primiparous rats only (IL-4: r=0.8759, p=0.002; IL-6: r=0.9206, p<0.001; IL10: r=0.8075, p=0.008; IL-13: r=0.8049, p=0.009; and while there were positive correlations in other cytokines, they did not survive Bonferroni correction including: IL-β: r=0.7497, p=0.02; IFNγ: r= 0.7596, p=0.018; IL-5: r=0.7452, p=0.021) but no significant correlations with KC/GRO or TNFα (p’s >0.27). There were two positive correlations in nulliparous rats given Premarin but these do not survive Bonferroni correction (KC/GRO: r=0.7717, p=0.015 and IL-13: r=0.6838, p=0.042). Primiparous rats given oil had two positive correlations with IL-10 and IL-5 and BrdU-labelled cells (r=0.8341, p=0.01; r=0.8291, p=0.011). There were no significant correlations among cytokines and neurogenesis for nulliparous oil-treated rats (p’s >0.36). There were no significant correlations with the probe trial and cytokines for any group or for levels of estrogens except for in the nulliparous group given Premarin with IL-4 (r=0.8746, p=0.004).

Negative correlations with cytokines and learning were seen in nulliparous rats treated with oil in the early learning (days 1-3) days (KC/GRO: r=-0.71, p=0.00202; TNF-α: r=-0.88, p=0.001) but these correlations do not reach significance in the Premarin treated groups (p=0.17, 0.11, respectively).

There was a positive correlation between ventral DCX-expression and distance travelled to reach the hidden platform on days 1 and 2 of training in the Premarin-treated primiparous rats only (r=0.7389, p<0.0001) but a non-significant negative correlation in nulliparous rats.

## Discussion

Parity influenced learning, neurogenesis and response to the hormone therapy, Premarin, in middle-aged female rodents. In middle age, primiparity was associated with more new neurons in the dentate gyrus, a greater proportion of proliferative immature neurons in the dentate gyrus, and a greater expression of zif268 in the CA3 region compared to nulliparity. Furthermore, previously parous rats exhibited more irregular estrous cycling, higher levels of serum estrone compared to nulliparous controls in middle age. Premarin impaired early spatial learning, reversal learning (trial one) and decreased immature neurons in the ventral dentate gyrus in primiparous rats, while Premarin had a beneficial effect on learning in nulliparous rats. Some of the effects also differed by ovarian hormone status, as long-term ovariectomy enhanced spatial memory and reduced zif268 expression in the CA1 and CA3 region. Furthermore, Premarin increased zif268 expression in the ventral CA3 region and increased adrenal to body mass ratio in primiparous, but not nulliparous rats. However, nulliparous rats had higher serum levels of the cytokines TNFα and KC/GRO with Premarin compared to primiparous rats. These findings indicate that parity has long-lasting effects on brain, immune signalling and behaviour in middle-aged females, and that reproductive history alters the ability of the hormone therapy, Premarin, to exert its effects on neural and cognitive effects in mid-life.

### Premarin inhibited early spatial learning and reversal learning in primiparous rats but enhanced learning and memory in nulliparous rats

In the present study, Premarin impaired early spatial training in primiparous but enhanced early spatial training in nulliparous animals. In addition, we found the same relationship between parity and Premarin on early reversal training with Premarin impairing performance in primiparous rats only. Furthermore, we found that Premarin enhanced memory in intact nulliparous, but not primiparous, rats. These findings are consistent with other studies that show that the same dose of Premarin enhanced working and reference memory in nulliparous middle-aged rats (Acosta et al. 2009; Acosta et al., 2010). These same effects were not seen in young adult nulliparous females as another study from our laboratory showed that lower doses of Premarin impaired both reference and working memory in the radial arm maze in young adult nulliparous rats (Barha and Galea, 2013). Collectively, these results suggest that higher doses of Premarin facilitate learning and memory in middle-aged nulliparous rats, but intriguingly, our novel data suggests Premarin impairs learning and memory in middle-aged primiparous rats. These findings also shed light on the finding from meta-analyses that indicate little cognitive benefit from Premarin in postmenopausal women (Ryan et al., 2008; Hogevorst et al., 2000). Because the majority of women are parous, the impairments (or lack of benefit) may be due to the effects of parity on Premarin’s ability to influence spatial and reversal learning.

In the present study, ovarian status interacted with some of the findings with long-term ovariectomy improving spatial memory in primiparous, but not nulliparous, rats. However, in another study, short-term ovariectomy impaired spatial memory in middle-aged multiparous rats (Barha et al., 2015). Another study compared long-term versus short term ovariectomy in much older female rats (~22 months) and found that long-term ovariectomy improved spatial memory in nulliparous rats (Bimonte-Nelson et al., 2003), indicating age and time since ovariectomy may play a role in how ovariectomy influences memory. In the current study, long-term ovariectomy only significantly improved spatial memory in primiparous rats but the means favored the same outcome (long-term ovariectomy improving memory) in nulliparous rats, consistent with previous work in older rats (Bimonte-Nelson et al., 2003). Collectively, these findings suggest that the duration of ovarian hormone deprivation can influence spatial memory, with short-term ovariectomy impairing, and long-term ovariectomy improving, spatial memory, regardless of parity.

### Primiparity is associated with greater neurogenesis levels in the dentate gyrus, while Premarin reduces neurogenesis in both nulliparous and primiparous rats dependent on region and age of new neuron

In middle-aged females, primiparity was associated with increased neurogenesis (BrdU-labelled cells) compared to nulliparity. This is consistent with another study from our group indicating that multiparous middle-aged females had higher levels of immature neurons (Barha et al., 2015). Curiously, in this study we only saw a greater number of immature neurons (DCX) in the primiparous animals in the ventral dentate gyrus while in multiparous rats a previous study showed an increased in DCX-expression when both dorsal and ventral dentate gyrus counts were combined (Barha et al., 2015). This slight difference may be due to the amount of parity, as the Barha study used retired breeders who had at least 4 litters while in this experiment, primiparous animals were used. Indeed, we have previously shown that the amount of parity affects neurogenesis in the early postpartum, with primiparous, but not biparous rats, showing a reduction in the survival of BrdU-labelled cells (labelled in the early postpartum and surviving until late postpartum; Pawluski and Galea, 2007). Amount of parity also affects verbal memory deficits during pregnancy, with multigravid women having worse scores than primigravid women (Glynn, 2012). Taken together, these findings suggest that parity influences neurogenesis in the early postpartum and well into middle-age.

In the present study Premarin treatment increased the survival of BrdU-labelled cells in the dorsal dentate gyrus which mirrors what is seen in young adult nulliparous females (Barha and Galea, 2013). Primiparous rat also had greater proportion of type 1 proliferative neurons, which may indicate that the enhanced neurogenesis with parity is associated with increased cell proliferation. It is also possible that parity influences the maturation of new neurons in the hippocampus that may explain some of the differences detected in the present study. However, we also found that Premarin reduced neurogenesis (BrdU-labelled cells) in the ventral dentate gyrus dramatically in the sham-operated animals relative to OVX counterparts. These findings are in contrast to what was seen previously in young adult nulliparous ovariectomized rats, where chronic 34 days of Premarin increased neurogenesis (Barha and Galea, 2013). However, there are a number of differences between the two studies that can explain this disparity in findings. First, BrdU was given prior to Premarin in the earlier study (Barha and Galea, 2012), unlike in the present study, where BrdU was given on day 2 after Premarin treatment. When BrdU is administered prior to any treatment, this will examine the effects of Premarin to influence survival of new neurons independent of any possible change in cell proliferation. In the present study, giving BrdU after the initial treatment with Premarin determined the influence of Premarin to produce and maintain new neurons under a Premarin-rich environment. Importantly, in the 2013 study, Premarin was given to *young* adult nulliparous OVX rats, and neurogenesis assessment was not divided into dorsal versus ventral regions of the hippocampus.

The Premarin-induced reduction in neurogenesis is also evident when examining immature neurons (DCX-expressing cells), but with regional differences as the reduction is seen in the ventral dentate gyrus of primiparous rats and the dorsal dentate gyrus of nulliparous rats. However, as described earlier Premarin reduced more mature neurons (the BrdU-labelled cells) in the ventral region, regardless of parity. Although it may seem puzzling that we see regional differences between number of DCX-expressing cells versus BrdU-labelled cells with parity and Premarin, this is likely due to the broad spread of ages of immature neurons in DCX-expressing cells, given that DCX is expressed for 21 days in rats (Brown et al., 2003). That is, doublecortin is expressed in all immature neurons ranging from hours to 21-30 days of age, whereas BrdU-labelled cells in this study were three-week old cells that had been synthesizing DNA for a 2 h period 21 days prior to termination. It is also important to keep in mind that new neurons may not function similarly in nulliparous versus primiparous animals. Indeed, we found more immature neurons in the ventral dentate were associated with poorer in the Premarin-treated primiparous, but not nulliparous, rats. These findings suggest that the relationship between neurogenesis and acquisition depends on previous parity when under the influence of Premarin.

### Primiparous rats had greater activation (zif268 expression) in the CA3 region, while treatment with Premarin increased expression in the ventral CA3 in primiparous rats only

Primiparous rats exhibited increased zif268 expression in the CA3 region. This suggests that neuronal activation levels were higher in primiparous compared to nulliparous rats in response to spatial memory retrieval. This is partially consistent with findings of greater LTP, BDNF and enhanced pCREB in older multiparous mice compared to nulliparous mice (Tomizawa et al., 2003). Intriguingly, the same increase in zif268 expression was also seen in young adult nulliparous rats after different types of hippocampus-dependent learning compared to males (Yagi et al., 2015, 2017), suggesting that activation of the CA3 region is particularly important for cognition in female rats. Given that we saw increased CA3 activation in the primiparous rats only this may suggest that primiparous rats have activation patterns similar to young adult female rats after spatial learning. This is reminiscent of the finding that estrogens upregulated cell proliferation in multiparous middle-aged and nulliparous young adult rats, but not nulliparous middle-aged rats (Barha and Galea, 2011). Collectively these studies indicate that parity may protect the hippocampal response to aging. However, future studies need to determine whether these cell-signalling and neurotrophic factors are responsible for the differences in response to hormone therapy in older females dependent on reproductive history.

Neither Premarin nor parity altered activation of new neurons in response to spatial memory retrieval. This is in contrast to the study in young nulliparous rats given a lower dose of Premarin that saw decreased activation in a different task (Barha and Galea, 2013). The coexpression of BrdU/zif268 was very low in the present study, likely due to the older age of the rats compared to the previous study in young adult rats. Intriguingly, however, Premarin increased activation of neurons in the CA3 region of the hippocampus in response to spatial memory retrieval but only in the primiparous rats. Because Premarin impaired spatial performance and reversal learning this suggests that the increased activation of CA3 neurons may have interfered with performance in primiparous rats. Further research should examine the relationship and functioning of new neurons under different treatments with differing reproductive history.

### Possible mechanisms of long-lasting influence of reproductive experience on neurogenesis and spatial learning: immune signalling, estrogens, and adrenal steroids

TNF-α and KC/GRO were increased by Premarin in nulliparous rats but not in primiparous rats. Interestingly, TNF-α has beneficial effects on synaptic strengthening (Beattie et al., 2002) and learning and memory (Baune et al., 2008) in the healthy adult brain. Thus, the effect of Premarin to enhance learning in nulliparous rats may be related to its effect to increase TNF-α. Further, the same relationship of enhanced KC/GRO with aging and improved cognition is seen in aged male rats, as KC/GRO is also upregulated in the hippocampus after running (Speisman et al., 2013), suggesting that the enhancement in cognition may be related to increased KC/GRO in the present study. Indeed, better performance on the first three days of training was related to both increased KC/GRO and TNFα in the oil-treated Nulliparous, but not Primiparous, groups. These results suggest differing relationships of circulating cytokine levels to learning in nulliparous dependent on previous parity.

Previous data from our own laboratory in middle-aged rats indicate differences in cytokine levels dependent on previous parity, in which primiparous rats have lower serum IFNγ, IL-10 and IL-4 compared to their nulliparous counterparts (Mahmoud et al., under revision). However, in the present study we did not see significant changes in these cytokines, perhaps due to the fact that they had also undergone cognitive testing in this study.

Intriguingly, several cytokines (Il-6, IL-4, IL-10, IL-13) were positively associated with neurogenesis in primiparous rats treated with Premarin, but the same significant correlations were not seen in nulliparous rats with or without Premarin. These data are intriguing but also somewhat consistent with the findings of Urza et al 2017, showing that only after a challenge (ovarian tumor) did cytokines respond differentially in previous parous versus nulliparous mice. In our study, while we did not have an immune challenge, the rats did undergo cognitive testing, surgeries, and injections. These findings suggest that previous parity interacts with aging changing the relationship of neurogenesis with cytokine levels after hormone treatment.

Premarin enhanced adrenal to body mass ratio in primiparous rats only which suggests that Premarin increases CORT in primiparous but not nulliparous, rats, as adrenal mass has been linked to serum CORT (Chan et al., 2014). As high CORT can interfere with learning and memory (Conrad, 2010) it is possible that Premarin worked via increased adrenal steroids to negatively affect learning in primiparous compared to nulliparous rats. The question of why this occurred is an open one, although primiparous rats had higher levels of serum estrone concentrations, were more likely to cycle irregularly, but no significant differences in serum estradiol. This later finding is interesting because estradiol levels show a dose-dependent relationship with spatial working memory with lower levels facilitating performance compared to absent or very high levels of estradiol (Holmes et al., 2002). However, estrone levels are generally associated with poorer memory in young adult nulliparous rats (Barha et al., 2009). Thus, it is possible that higher levels of estrone or estrone/estradiol ratio in primiparous rats contributed to slightly better learning in the Morris water maze. Collectively these data suggest that levels of adrenal mass and specific cytokines are altered with previous parity and with exposure to Premarin. More studies are needed to identify neuroimmune interactions on cognition with parity and aging.

## Conclusions

The present study shows that previous reproductive experience can increase neurogenesis, spatial acquisition, estrone levels and disrupt cycling in middle-aged rats. Intriguingly, treatment with Premarin, the CEE hormone therapy, differentially affected middle-aged rats, dependent on previous reproductive experience. Premarin negatively influenced spatial acquisition while counterintuitively increasing activation of neurons (zif268) in the ventral CA3 region in primiparous rats only, suggesting that previous parity also interacts with hormone therapy to exert effects on cognitive and neural measures. Premarin also increased serum cytokines TNFα and KC/GRO in nulliparous but not primiparous rats. These findings add to the growing field of how reproductive experience can have long-lasting influences on the physiology of the middle-aged female, but also suggest that parity may need to be taken into account when considering the influence of hormone therapies in older age. The fact that previous parity influences the efficacy of hormone treatment contributes to a growing research indicating that other treatments such as selective serotonin reuptake inhibitors influence coping behavior and neurogenesis differently depending on parity (Workman et al., 2016; Gobinath et al., 2017; Overgaard et al., in press). Clearly, we are in the infancy of this field of determining how maternal experience influence the aging brain. In particular, there are clinical implications that women may need to consider reproductive history in their selection of HT and assessment of risk versus benefit of HT use to combat cognitive menopausal symptoms. In summary, our current data suggests that Premarin resulted in different, and sometimes opposing, outcomes on brain and behaviour in middle-aged rats dependent on parity. These findings underscore the necessity of considering reproductive history in individualized treatments in ageing women.

## Acknowledgements

This work was funded by a CIHR (PJT148662) and Alzheimer Society of Canada operating grants to LAMG.

## References

Acosta JI, Mayer LP, Braden BB, Nonnenmacher S, Mennenga SE, Bimonte-Nelson HA. The cognitive effects of conjugated equine estrogens depend on whether menopause etiology is transitional or surgical. Endocrinology 2010;151:3795-804. doi: 10.1210/en.2010-0055. Epub 2010 Jun 16.

Acosta JI, Mayer L, Talboom JS, Zay C, Scheldrup M, Castillo J, Demers LM, Enders CK, Bimonte-Nelson HA. Premarin improves memory, prevents scopolamine-induced amnesia and increases number of basal forebrain choline acetyltransferase positive cells in middle-aged surgically menopausal rats. Horm Behav. 2009;55:454-64. doi: 10.1016/j.yhbeh.2008.11.008. Epub 2008 Dec 3.

Barha CK, Galea LA. The hormone therapy, Premarin, impairs hippocampus-dependent spatial learning and memory and reduces activation of new granule neurons in response to memory in female rats. Neurobiol Aging. 2013;34:986-1004. doi: 0.1016/j.neurobiolaging.2012.07.009.

Barha CK, Lieblich SE, Chow C, Galea LA Multiparity-induced enhancement of hippocampal neurogenesis and spatial memory depends on ovarian hormone status in middle age. Neurobiol Aging. 2015;36:2391-405. doi: 10.1016/j.neurobiolaging.2015.04.007

Barrat F, Lesourd B, Boulouis HJ, Thibault D, Vincent-Naulleau S, Gjata B, Louise A, Neway T, Pilet C. Sex and parity modulate cytokine production during murine ageing. Clin Exp Immunol. 1997;109:562-8.

Barrett ES, Parlett LE, Windham GC, Swan SH. Differences in ovarian hormones in relation to parity and time since last birth. Fertil Steril. 2014;101:1773-80.e1

Baune BT, Wiede F, Braun A, Golledge J, Arolt V, Koerner H. Cognitive dysfunction in mice deficient for TNF- and its receptors. Am J Med Genet B Neuropsychiatr Genet. 2008;147B:1056-64. doi: 10.1002/ajmg.b.30712.

Beattie EC, Stellwagen D, Morishita W, Bresnahan JC, Ha BK, Von Zastrow M, Beattie MS, Malenka RC. Control of synaptic strength by glial TNFalpha. Science. 2002;295:2282-5.

Beeri MS, Rapp M, Schmeidler J, Reichenberg A, Purohit DP, Perl DP, Grossman HT, Prohovnik I, Haroutunian V, Silverman JM. Number of children is associated with neuropathology of Alzheimer's disease in women. Neurobiol Aging. 2009;30:1184-91.

Bridges RS, Byrnes EM. Reproductive experience reduces circulating 17beta-estradiol and prolactin levels during proestrus and alters estrogen sensitivity in female rats. Endocrinology. 2006;147:2575-82.

Cameron HA, Woolley CS, McEwen BS, Gould E. Differentiation of newly born neurons and glia in the dentate gyrus of the adult rat. Neuroscience. 1993;56:337-44.

Colucci M, Cammarata S, Assini A, Croce R, Clerici F, Novello C, Mazzella L, Dagnino N, Mariani C, Tanganelli P. The number of pregnancies is a risk factor for Alzheimer's disease. Eur J Neurol. 2006;13:1374-7.

Conrad, CD. A critical review of chronic stress effects on spatial learning and memory. Prog Neuropsychopharmacol Biol Psychiatry 2010; 34:742-755.

Cramer DW, Vitonis AF. Signatures of reproductive events on blood counts and biomarkers of inflammation: Implications for chronic disease risk. PLoS One. 2017;12:e0172530.

Cui J, Jothishankar B, He P, Staufenbiel M, Shen Y, Li R. Amyloid precursor protein mutation disrupts reproductive experience-enhanced normal cognitive development in a mouse model of Alzheimer's disease. Mol Neurobiol. 2014; 49:103-12.

Engler-Chiurazzi E, Tsang C, Nonnenmacher S, Liang WS, Corneveaux JJ, Prokai L, Huentelman MJ, Bimonte-Nelson HA. Tonic Premarin dose-dependently enhances memory, affects neurotrophin protein levels and alters gene expression in middle-aged rats. Neurobiol Aging. 2011 Apr;32(4):680-97. doi: 10.1016/j.neurobiolaging.2009.09.005.

Fox M, Berzuini C, Knapp LA. Cumulative estrogen exposure, number of menstrual cycles, and Alzheimer's risk in a cohort of British women. Psychoneuroendocrinology. 2013;38:2973-82.

Galea LA, Leuner B, Slattery DA. Hippocampal plasticity during the peripartum period: influence of sex steroids, stress and ageing. J Neuroendocrinol. 2014;26:641-8. doi: 10.1111/jne.12177.

Gatewood JD, Morgan MD, Eaton M, McNamara IM, Stevens LF, Macbeth AH, Meyer EA, Lomas LM, Kozub FJ, Lambert KG, Kinsley CH. Motherhood mitigates aging-related decrements in learning and memory and positively affects brain aging in the rat. Brain Res Bull. 2005;66:91-8.

Ghaebi M, Nouri M, Ghasemzadeh A, Farzadi L, Jadidi-Niaragh F, Ahmadi M, Yousefi M. Immune regulatory network in successful pregnancy and reproductive failures. Biomed Pharmacother. 2017;88:61-73. doi: 10.1016/j.biopha.2017.01.016.

Glynn LM. Increasing parity is associated with cumulative effects on memory. J Womens Health (Larchmt). 2012;21:1038-45. doi: 10.1089/jwh.2011.3206.

Gibbs RB. Estrogen therapy and cognition: a review of the cholinergic hypothesis. Endocr. Rev. 2010; 31:224-253.

Grindheim G, Toska K, Estensen ME, Rosseland LA. Changes in pulmonary function during pregnancy: a longitudinal cohort study. BJOG. 2012;119:94-101.

Heys M, Jiang C, Cheng KK, Zhang W, Au Yeung SL, Lam TH, Leung GM, Schooling CM. Life long endogenous estrogen exposure and later adulthood cognitive function in a population of naturally postmenopausal women from Southern China: the Guangzhou Biobank Cohort Study. Psychoneuroendocrinology. 2011;36:864-73. doi: 10.1016/j.psyneuen.2010.11.009.

Hoekzema E, Barba-Müller E, Pozzobon C, Picado M, Lucco F, García-García D, Soliva JC, Tobeña A, Desco M, Crone EA, Ballesteros A, Carmona S, Vilarroya O. Pregnancy leads to long-lasting changes in human brain structure. Nat Neurosci. 2017;20:287-296. doi: 10.103 8/nn.4458.

Hogervorst E, Williams J, Budge M, Riedel W, Jolles J. The nature of the effect of female gonadal hormone replacement therapy on cognitive function in post-menopausal women: a metaanalysis. Neuroscience. 2000; 101:485-512.

Holl K, Lundin E, Kaasila M, Grankvist K, Afanasyeva Y, Hallmans G, Lehtinen M, Pukkala E, Surcel HM, Toniolo P, Zeleniuch-Jacquotte A, Koskela P, Lukanova A. Effect of long-term storage on hormone measurements in samples from pregnant women: The experience of the Finnish Maternity Cohort Acta Oncologica 2008;47:406-12.

Imtiaz B, Taipale H, Tanskanen A, Tiihonen M, Kivipelto M, Heikkinen AM, Tiihonen J, Soininen H, Hartikainen S, Tolppanen AM. Risk of Alzheimer's disease among users of postmenopausal hormone therapy: A nationwide case-control study. Maturitas. 2017;98:7-13.

Karim R, Dang H, Henderson VW, Hodis HN, St John J, Brinton RD, Mack WJ. Effect of Reproductive History and Exogenous Hormone Use on Cognitive Function in Mid- and Late Life. J Am Geriatr Soc. 2016;64:2448-2456.

Lemaire V, Billard JM, Dutar P, George O, Piazza PV, Epelbaum J, Le Moal M, Mayo W. Motherhood-induced memory improvement persists across lifespan in rats but is abolished by a gestational stress. Eur J Neurosci. 2006;23:3368-74.

Love G, Torrey N, McNamara I, Morgan M, Banks M, Hester NW, Glasper ER, Devries AC, Kinsley CH, Lambert KG. Maternal experience produces long-lasting behavioral modifications in the rat. Behav Neurosci. 2005;119:1084-96.

Macbeth AH, Scharfman HE, Maclusky NJ, Gautreaux C, Luine VN. Effects of multiparity on recognition memory, monoaminergic neurotransmitters, and brain-derived neurotrophic factor (BDNF). Horm Behav. 2008; 54:7-17.

McClure RE, Barha CK, Galea LA. 17ß-Estradiol, but not estrone, increases the survival and activation of new neurons in the hippocampus in response to spatial memory in adult female rats. Horm Behav. 2013;63:144-57. doi: 10.1016/j.yhbeh.2012.09.011.

Parsons CE, Young KS, Petersen MV, Jegindoe Elmholdt EM, Vuust P, Stein A, Kringelbach ML. Duration of motherhood has incremental effects on mothers' neural processing of infant vocal cues: a neuroimaging study of women. Sci Rep. 2017;7:1727. doi: 10.1038/s41598-017-01776-3.

Pawluski JL, Galea LA. Hippocampal morphology is differentially affected by reproductive experience in the mother. J Neurobiol. 2006;66:71-81.

Pawluski JL, Vanderbyl BL, Ragan K, Galea LA. First reproductive experience persistently affects spatial reference and working memory in the mother and these effects are not due to pregnancy or 'mothering' alone. Behav Brain Res. 2006;175:157-65.

Ptok U, Barkow K, Heun R. Fertility and number of children in patients with Alzheimer's disease. Arch Womens Ment Health. 2002;5:83-6.

Roes, M.M., Galea, L.A.M. (2015). The maternal brain: short and long-term effects of reproductive experience on hippocampus structure and function in adulthood. In R. Shansky & J. Johnson (Eds.) Sex Differences in the Central Nervous System, Elsevier. pp. 197-220

Rossetti MF, Varayoud J, Lazzarino GP, Luque EH, Ramos JG. Pregnancy and lactation differentially modify the transcriptional regulation of steroidogenic enzymes through DNA methylation mechanisms in the hippocampus of aged rats. Mol Cell Endocrinol. 2016;429:73-83

Savu O, Jurcuţ R, Giuşcä S, van Mieghem T, Gussi I, Popescu BA, Ginghinä C, Rademakers F, Deprest J, Voigt JU. Morphological and functional adaptation of the maternal heart during pregnancy. Circ Cardiovasc Imaging. 2012;5:289-97.

Speisman RB, Kumar A, Rani A, Foster TC, Ormerod BK. Daily exercise improves memory, stimulates hippocampal neurogenesis and modulates immune and neuroimmune cytokines in aging rats. Brain Beh Immun. 2013;28:25-43. doi:10.1016/j.bbi.2012.09.013.

Tomizawa K, Iga N, Lu YF, Moriwaki A, Matsushita M, Li ST, Miyamoto O, Itano T, Matsui H. Oxytocin improves long-lasting spatial memory during motherhood through MAP kinase cascade. Nat Neurosci. 2003;6:384-90.

Urzua U, Chacon C, Lizama L, Sarmiento S, Villalobos P, Kroxato B, Marcelain K, Gonzalez MJ. Parity History Determines a Systemic Inflammatory Response to Spread of Ovarian Cancer in Naturally Aged Mice. Aging Dis. 2017;8:546-557

Walf AA, Paris JJ, Frye CA. Chronic estradiol replacement to aged female rats reduces anxiety-like and depression-like behavior and enhances cognitive performance. Psychoneuroendo 2009;34:909-916.

Woolley CS, Gould E, Frankfurt M, McEwen BS. Naturally occurring fluctuation in dendritic spine density on adult hippocampal pyramidal neurons. J Neurosci. 1990;10:4035-9.

